# Sex differences in central inflammatory pain sensitization are associated with differential expression of glycine receptors and GLP-1 at the spinal cord

**DOI:** 10.1101/2019.12.31.892133

**Authors:** TA. Mariqueo, G. Améstica, J. Pino, R. Barra, J. Stehberg, W. González, K. Castillo

**Author notes:** Author responsible for correspondence: Trinidad A. Mariqueo, Center for Medical Research (CIM), Laboratory of Neuropharmacology, School of Medicine, Universidad de Talca, Talca, Chile Av. Lircay s/n, Talca 3460000, Chile.

## Abstract

**Background:** Females have higher inflammatory pain representation. However, sex differences in central pain sensitization and the regulation of nociceptive response to peripheral inflammation remain unclear. The central pain sensitization is mediated by inhibitory neurotransmission and glial cell activity dysregulation where spinal glycine and GLP-1 receptors have described play a critical role.

**Objectives:** The aim of this study was to compare the mechanical withdrawal nociceptive threshold with spinal glycine receptor subunits and GLP-1 expression in adult male and female rats after inflammatory hypersensitivity.

**Methods:** Sex differences in inflammatory nociception were evaluated before and after intraplantar hindpaw Zymosan A injection in Sprague-Dawley rats. Mechanical paw withdrawal thresholds were tested using von Frey filaments.

Western blot was used to measure GlyRs subunits protein levels in the spinal cord. GLP-1 was determined using the Magnetic Luminex Assay.

**Results:** A reduced nociceptive threshold was observed in males and females rats after 4 hours of inflammatory Zymosan A injection. However, this reduction was significantly major in females. Western blot analysis demonstrated significantly increased α1, α2, α3 and β GlyR subunit levels in male rats. Female rats only increased α3 and β GlyR subunits after Zymosan A injection. GLP-1 was reduced in female spinal tissues after an inflammatory injury.

**Conclusions:** Our study indicates that sex differences in nociceptive threshold after inflammatory Zymosan A rat pain sensitization is related to the sex differences in glycine receptor subunits and GLP-1 expression at the spinal cord.

## Background

Chronic pain constitutes a major health challenge, affecting millions of people Worldwide. At least 116 million of adults are affected in the United States alone, representing an important medical and economic burden to the Society and the families of the individuals affected [1].

The neural processing in the pathological pain is particularly associated with maladaptive spinal endogenous compensatory mechanisms [2]. The imbalance between excitatory neurotransmission mediated by glutamatergic N-methyl-D-aspartate (NMDA) and α-amino-3-hydroxy-5-methyl-4-isoxazolepropionic acid (AMPA) Receptors and inhibitory neurotransmission exerted by Gamma-aminobutyric acid (GABAA) and Glycine Receptors (GlyRs) play a critical role in the central pain sensitization process [3]. The activation of glutamate NMDA receptors triggers calcium-dependent signaling pathways, promoting the plasticity of central sensory processing circuits [4]. In addition, inflammation leads to the local synthesis and release of prostaglandin 2 (PGE2) which activates PGE2 receptors inducing the phosphorylation of α3GlyR, leading to GlyR inhibition. GlyR inhibition in turn decreases spinal inhibition, contributing to inflammatory hypersensitivity. Hence, mice deficient in PG2R and α3GlyR subunit show a reduced inflammatory hypersensitivity [5].

Persistent evidence indicates a higher prevalence of inflammatory pain among female patients [6,7]. The sex-hormone regulation may account in part for these nociceptive differences. Indeed, female Dorsal Root Ganglion (DRG) neurons showed about two-time higher potentiation of the NMDA currents by 1, 3, 5 (10) -estratrieno-3, 17β-diol (17β-Estradiol) [8]. In contrast, it was reported that glycine-evoked currents were reversibly reduced by 17β-Estradiol in hippocampal and spinal neurons in culture [9]. The sex differences in the central pain sensitization process have not been studied in detail. This process involves spinal microglia and astrocytes release of pro and anti-inflammatory mediators [10]. Hence, a recent study has shown that microglial activation has a potent antinociceptive activity by a mechanism that involves the intestinal hormone Glucagon-like peptide-1 (GLP-1) receptor [11]. The activation of GLP-1R promotes the release of anti-inflammatory Interleukin 10 and β-endorphins at the spinal circuit [11].

Considering the relevance of GlyRs and GLP1 in the central inflammatory pain sensitization regulation, we evaluated the association of nociceptive sex differences with the expression of these molecules using an animal model of inflammatory pain.

## Methods

### Animals

Male and female Sprague-Dawley rats (250 g) were obtained from the Animal Facility of the University of Chile. Animal care and experimental protocols for this study were approved by the Institutional Committee of Bioethics of the University of Santiago of Chile (No.301) and followed the guidelines for ethical protocols and care of experimental animals established by the National Institute of Health, MD, USA and the International Association for the Study of Pain in conscious animals. Every effort was made to minimize animal suffering. Animals were bred and housed in controlled laboratory conditions, received standard rat chow diet and water ad libitum and were housed on a 12-h light/dark cycle at a constant room temperature of 23°C.

### Experimental procedures

The mechanical threshold measurements were performed using von Frey filaments by two different researchers who were blinded to the sexes of the animals. First, a group of 17 animals was used to evaluate the basal mechanical threshold of male and female rats. Then, a different group of male (n=6) and female (n=6) rats were evaluated 24 hours before and 4 hours after a right hindpaw Zymosan A injection. Immediately after the last evaluation, the spinal cord samples were obtained for Western blots and Luminex assays.

### Behavioral measurement

Rats were placed into individual chambers with wire mesh floors and transparent covers. After 15 minutes of habituation, their mechanical threshold was evaluated using the von Frey filaments manually. Perpendicular force was applied to the intraplantar zone of each paw using filaments with a constant pre-determined force (32-512 mN) from thinner to thicker until there was a withdrawal response. The mechanical withdrawal threshold was determined using the Up-Down method with 3 up-down cycles per measure [12].

### Inflammatory model of Pain

Inflammatory pain was induced by plantar right hindpaw injection of Zymosan A (from *Saccharomyces cerevisiae*, Sigma-Aldrich), 0.6 mg dissolved in 100 μL of saline solution (0.9% NaCl). The mechanical withdrawal threshold was determined 4 hours after Zymosan A injection in separated groups of male and female rats.

### Western Blotting

The left and right sides of the spinal cord (at spinal lumbar segments L5/6) were separated from the median sulcus using a surgical blade slide. The contralateral side from the Zymosan A injection was used as a control. Spinal cord tissue (60 mg) was homogenized in 500 μL of RIPA lysis buffer containing: 50 mM Tris-HCl, 150 mM NaCl; 1.0% (v/v) NP-40, 0.5% (w/v) Sodium Deoxycholate, 1mM EDTA, 0.1%(w/v) SDS dissolved in deionized water, pH of 7.4, and supplemented with a complete protease inhibitor cocktail (Pierce, Protease Inhibitor Mini Tablets, EDTA-Free). The samples were homogenized and centrifuged at 12000 g for 10 min. Total proteins from the supernatant were quantified using a Bradford Protein Assay Kit (Merck) and subjected to electrophoresis on 15% SDS–PAGE gels. Proteins were blotted onto nitrocellulose membranes (Thermo scientific) and blocked with 25 ml of 5% (w/v) skim milk in 1x Phosphate Buffered Saline with 0.05 % (v/v) Tween 20 (PBST) for 45 min with constant rocking at room temperature. Subsequently, the membranes were incubated with primary α1GlyR (1:500, mouse polyclonal IgG, Cat. No. 146111, Synaptic System) or α2GlyR (1:500, mouse monoclonal IgG, Cat. No. sc-398964, Santa Cruz Biotechnology) or α3GlyR (1:500, mouse polyclonal IgG, Cat. No. AB5472-50UL, Merck) or βGlyR (1:500, mouse monoclonal IgG, Cat. No. 146211, Synaptic System) subunits and GAPDH (1:2500, mouse monoclonal IgG, Cat. No. Sc-365062, Santa Cruz Biotechnology) antibodies dissolved in 5 ml of 5% (w/v) skim milk in PBST, overnight at 4°C. After 18 h, the membranes were washed three times in PBST and incubated 2 h with horseradish peroxidase (HRP) secondary antibodies (1:5000, Santa Cruz Biotechnology). This procedure was followed by three membrane washes with PBST. The immunoreactivity of the proteins was detected and visualized after incubating the membranes with SuperSignal West Femto Maximum Sensitivity Substrate (Thermo scientific). GAPDH was used as a loading control. The Western blot bands were quantified by using “ImageJ” software.

### Luminex bead-based multiplex assay

Concentration of GLP-1 was determined using the Magnetic Luminex Assay (R&D Systems Inc., Minneapolis, MN) and MAGPIX reader (Luminex Corporation, Austin, TX.). We used 50 μl of diluted sample (1: 2) in a 5-plex plate according to the manufacturer’s instructions.

### Data analysis

All data are presented as mean ± standard error of mean (SEM). Statistical analyses were performed using GraphPad Prism 6 software. One-way repeated measure analysis of variance (ANOVA) followed by the Bonferroni post hoc test were used for multiple comparisons of means in von Frey assays. One-way ANOVA followed by the Bonferroni post hoc was used for Western blots and Luminex assay measurements. *P-*value <0.05 was considered as significant.

## Results

### Baseline sex withdrawal threshold comparison

In order to explore possible sex differences in the mechanical withdrawal threshold, von Frey tests were performed in all paws of male and female rats. As shown in Figure 1, males showed a significantly reduced withdrawal of the mechanical threshold in the left forepaws (*P*<0.0001), whereas the females exhibited a significantly reduced withdrawal of the mechanical threshold in both forepaws in comparison with hindpaws (left paw *P*<0.0001; right paw *P*<0.05) (Fig. 1).

**Fig. 1:**
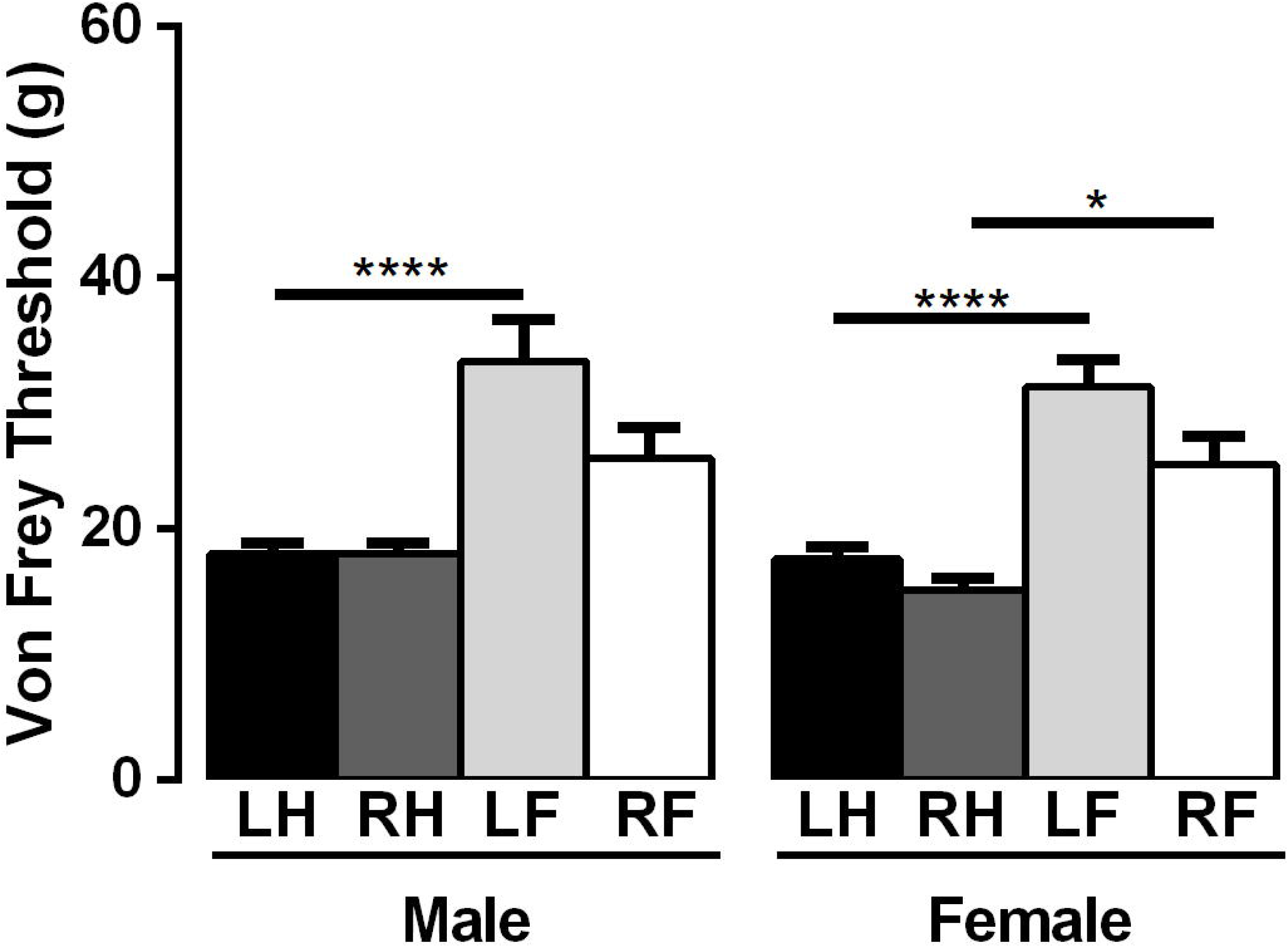
Behavioral evaluation of pain sensitization in the basal condition. Mechanical von Fray threshold was evaluated in all paws of male and female rats. Data are presented as mean ± SEM, (One-way ANOVA, Bonferroni Post Hoc Test, \**P*<0.05; \*\*\*\**P*<0.0001, n=8-9 in male and females, respectively). Abbreviations: left forepaw (LF), right forepaw (RF), left hindpaw (LH), right hindpaw (RH).

To determine sex differences in inflammatory pain, we induced paw inflammation by Zymosan A hindpaw injection. The inflammatory response was clearly visible after 4 hours (Fig. 2). Therefore, we evaluated the foot withdrawal to innocuous tactile stimulation before and after 4 hours of the Zymosan A injection. Although both sexes showed Zymosan A-induced allodynia, the response of females was higher than in males (males *P*<0.05; females *P*<0.0001) (Fig. 3). This evidence strongly supports the idea that females can perceive pain with a lower threshold compared to males, in a model of inflammatory pain.

**Fig. 2:**
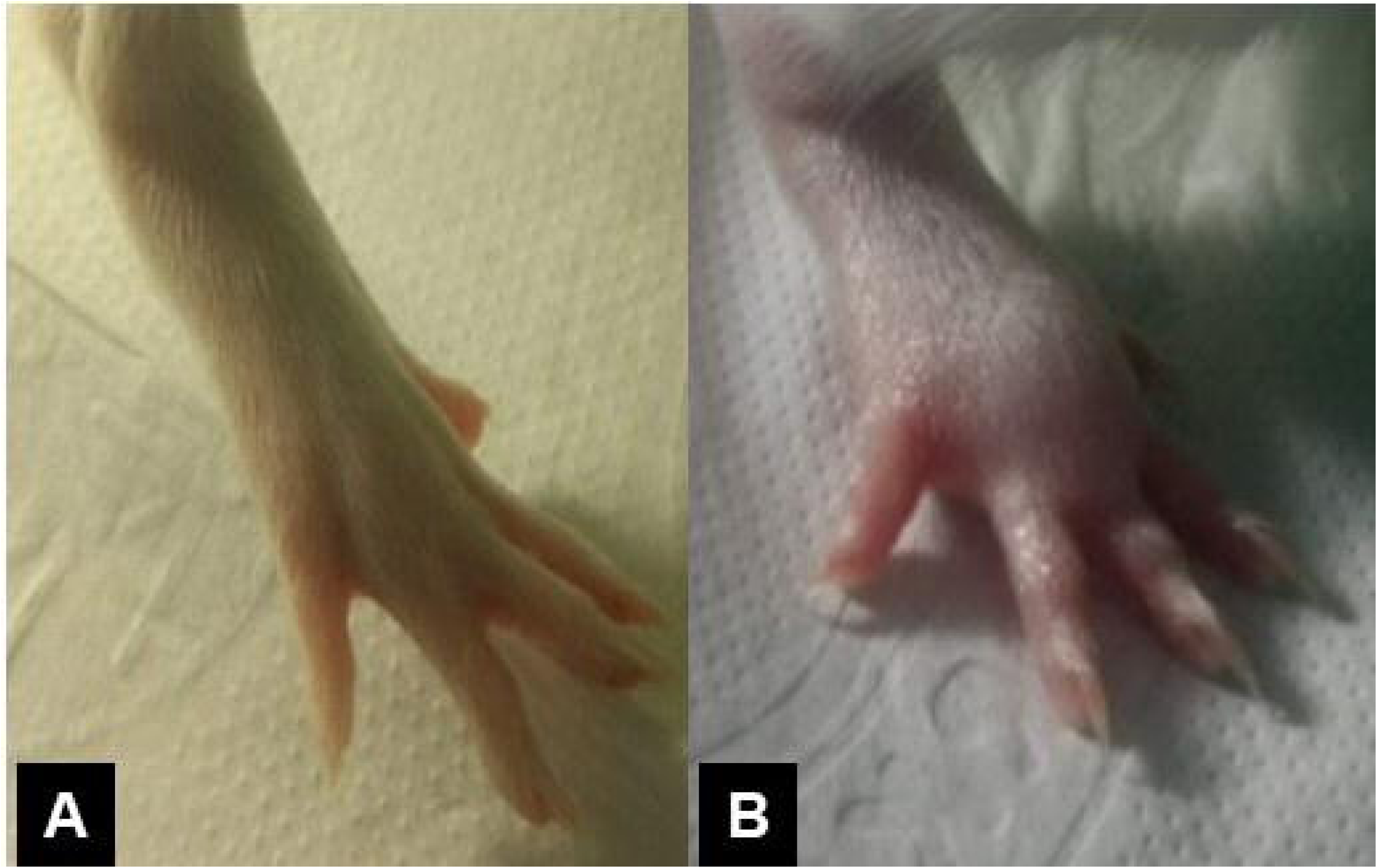
Zymosan A induced paw inflammation pain model. Representative images of a rat paw before (A) and after 4 hours of Zymosan A injection (B). Images are representative of 12 animals.

**Fig. 3:**
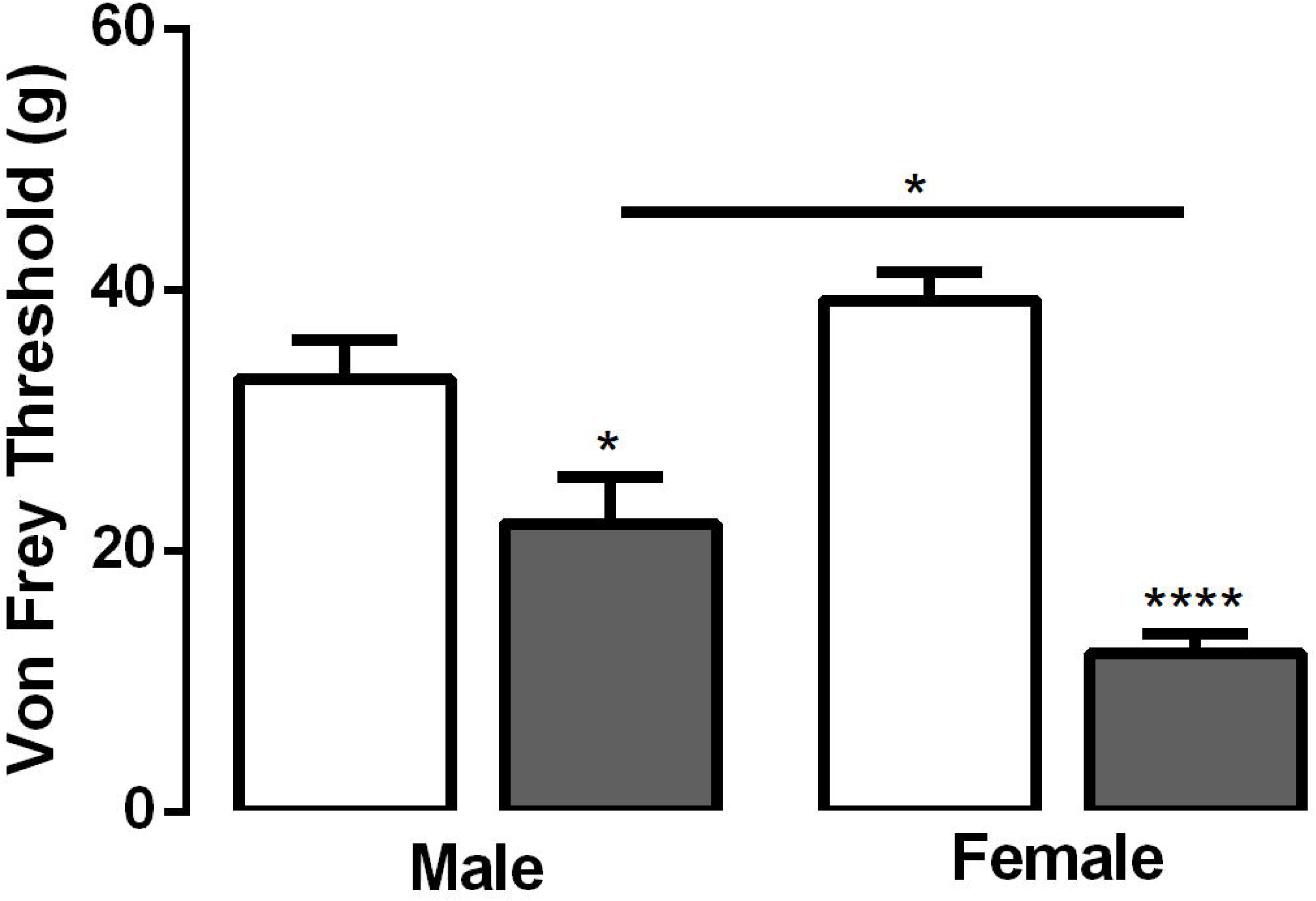
Mechanical von Frey nociceptive test evaluated in male and female rats before (white bars) and after 4 hours of Zymosan A injection (gray bars). Data are presented as mean ± SEM, *\*P*<0.05; *\*\*\*\*P*<0.0001 (One-way Anova, Bonferroni Post Hoc Test, n=12).

### Sex differences in central inflammatory pain sensitization

As GlyRs play a critical role in central inflammatory pain sensitization, we evaluated possible sex differences in GlyR subunit expression levels in spinal cord tissue 4 hours after a Zymosan A injection. Figure 4A-E shows the expression level of different GlyR subunits, analyzed by Western blot. The α1GlyR subunit increased in male spinal tissues after inflammatory stimulation (male *P*<0.001) (Fig. 4B). Similarly, the expression of the α2GlyR subunit appears to be increased only in male tissues (*P*<0.05) (Fig. 4C). Significant differences were observed in α3 subunit levels in both groups after the stimulus (male *P*<0.05; female *P*<0.001) (Fig. 4D). Comparable results were obtained for βGlyR subunit levels, exhibiting higher levels of β subunit after Zymosan A injection, in both male and females tissues (male *P*<0.05; female *P*<0.05) (Fig. 4E). These results suggest that increased inflammatory female sensitivity would be related in part with the lack of inhibitory compensation mechanisms at the spinal cord level. On the other hand, our data reveal the existence of fine tune mechanisms that regulate the GlyR subunit expression and modulation in the inflammatory pain context.

**Fig. 4:**
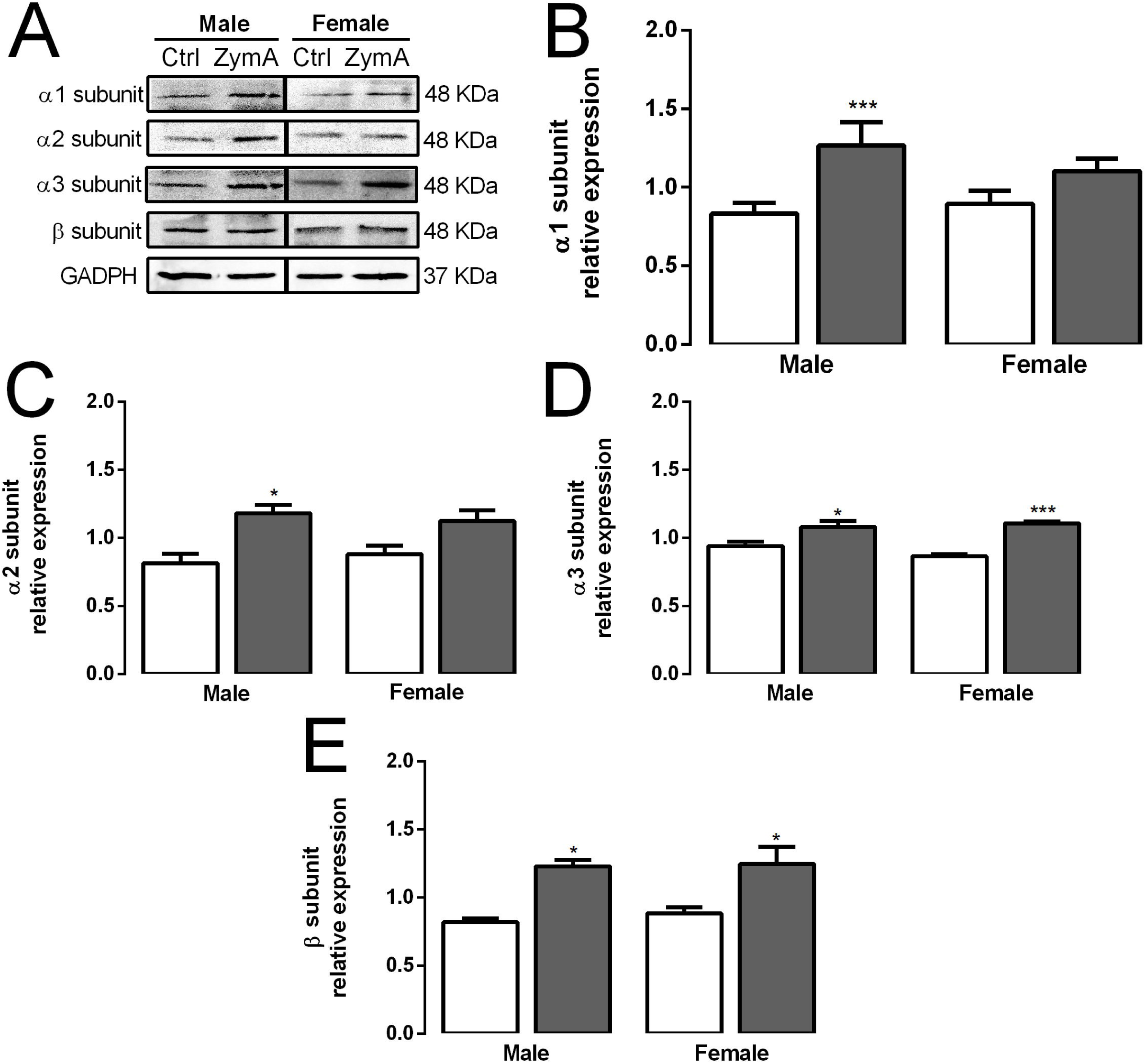
Expression analysis of different GlyR subunits levels in spinal cord tissues of inflammatory injured male and female rats. (A) Representative Western blots bands of the different GlyR subunits. (B-E) Comparison of GlyR subunits expression in male and female animals after Zymosan A injection (gray bars). Data are presented as mean ± SEM, \**P*<0.05; \*\*\**P* <0.001 (One-way ANOVA, Bonferroni Post Hoc Test, n= 8).

Previous studies demonstrated that GLP-1 receptor has a potent antinociceptive activity by a mechanism that involves microglial activation at the spinal cord level. Therefore, in the present study, we compare the expression of the GLP-1 hormone in male and female spinal cord derived tissues after 4 hours of a Zymosan injection. Interestingly, we found that males does not change GLP-1 in contrast to females which presented a significantly reduced GLP-1 expression after inflammatory injury (male *P*>0.05; female *P*<0.01) (Fig.5). Also, we found that basal level of GLP-1 was greater in females (*P*<0.01) (Fig.5). The microglial GLP-1 Receptors stimulate the anti-nociceptive response by the release of β-endorphins and anti-inflammatory interleukin 10 [13]. In this context, our results suggest that neuro-immune interface differences at spinal cord could be involved in the sex differences nociceptive responses, during inflammatory damage.

**Fig. 5:**
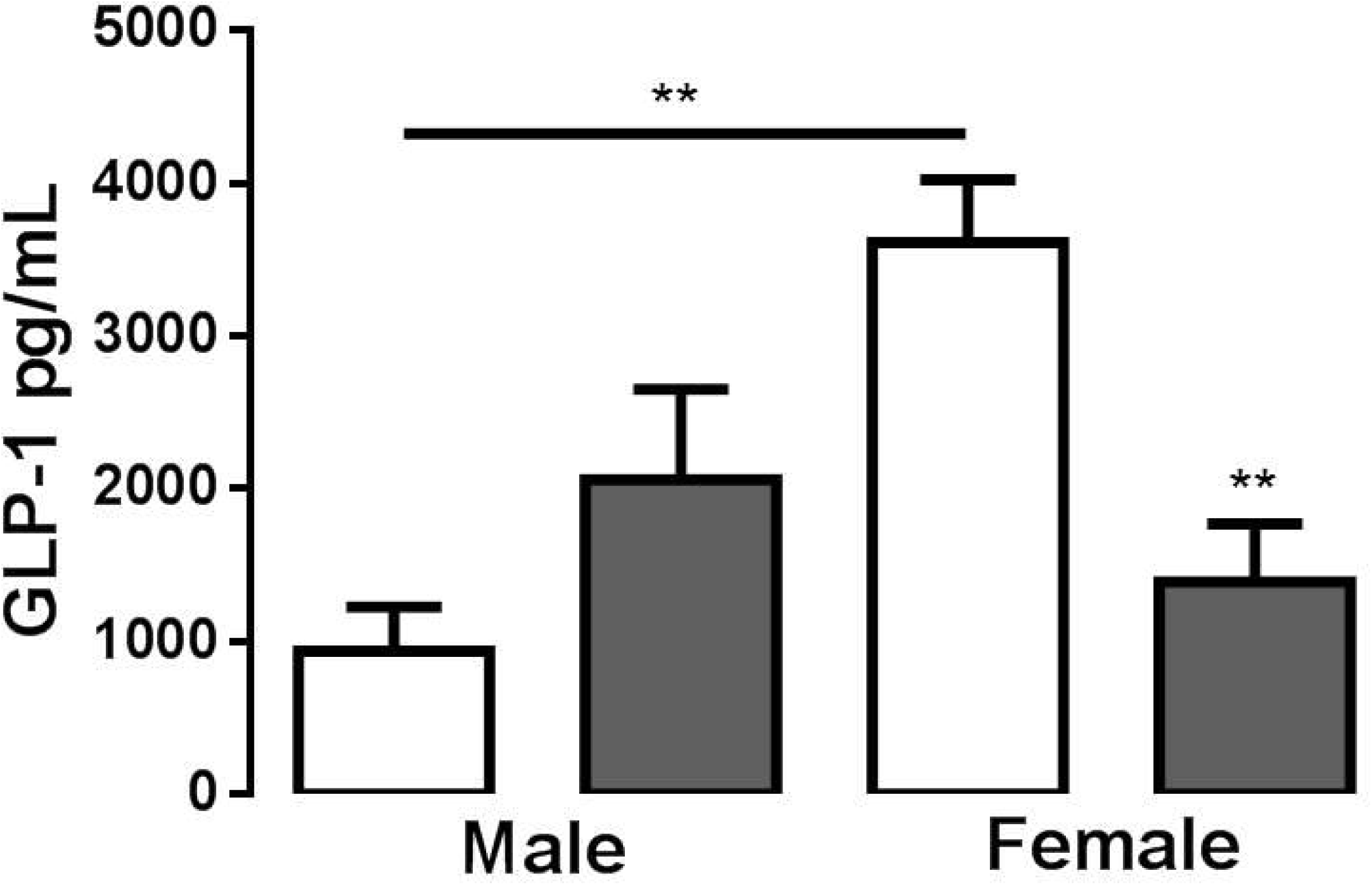
GLP-1 concentration (pg/mL) in spinal cord tissues derived from Zymosan A injected rats. GLP-1 before (white bars) and after 4 hours of Zymosan A injection (gray bars) in male and female rats. Data are presented as mean ± SEM, \**P*<0.05; \*\**P* <0.01 (One-way ANOVA, Bonferroni Post Hoc Test, n= 8).

## Discussion

In this study, we present evidence for sex differences in the withdrawal threshold and compensatory spinal inhibition mediated by GlyR and GLP-1 after peripheral pain induced by inflammatory Zymosan A injection. It is well known that acute peripheral nociceptive injury leads to chronic sensitization in central nociceptive circuits [4]. In this context, previous studies have shown higher prevalence and intensity of inflammatory pain in females [14]. Our results show that at basal levels males and females have similar withdrawal thresholds with a reduced response in forepaws in comparison with hindpaws. However, the nociceptive response after Zymosan A injection was greater in female than males, as measured by the Von Frey test. This sex difference in central pain sensitization could be attributed at least in part with hormonal pain modulation. Supportive of this statement it is the evidence that 17β-Estradiol modulates peripheral sensitization by increasing TRPV1 channel expression and peripheral sensitization of the primary afferent nociceptive neurons of female mice [15]. In the central pain sensitization, the modulation of NMDA, GABAA and Glycine Receptors plays a critical role. According with these evidences, it was shown that excitatory glutamatergic NMDAR-mediated currents were potentiated by 17β-Estradiol [8]. Consequently, this increase was found to be grater in female DRG neurons than in males (55±15% in female vs 19±7% in male) [8]. Furthermore, the GABAergic amplitude of miniature inhibitory postsynaptic currents (mIPSCs) in hippocampal slices was reduced by 17β-Estradiol through a mechanism involving a reduction of synaptic clustering of the Gephyrin scaffold protein [16]. Additionally, it was demonstrated that 17β-Estradiol inhibited glycine-evoked currents in cultured hippocampal neurons [9]. However, the sex differences in the expression of GlyR subunits in the context of inflammatory pain have not been explored, in spite of its clear importance in pain sensitivity.

Our results show that Zymosan A-induced inflammation is associated with an increased expression of spinal α1GlyR subunit in males. This result was in agreement with previously reported data that demonstrated that chronic CFA-induced inflammatory pain correlates with increased α1GlyR subunit expression [17]. The α1GlyR subunit is expressed in mature stages of central neural system development and exerts faster current parameters in comparison with α2 or α3 GlyR [18,19]. In addition, our results showed significant changes in the α2GlyR subunit expression. The α2GlyR have greater current conductance in comparison with the other types of homomeric GlyRs. Consistent with our data, Imlach et al., showed greater α2GlyR subunit expression and time constant of decay (representative of α2GlyR currents), in male spinal radial neurons after neuropathic pain [20]. Therefore, we propose that the increase in α2GlyR promotes the neural circuit hyperpolarization as a compensatory response in different types of pain. Furthermore, our results showed that both males and females had increased α3GlyR. This subunit is expressed in the dorsal horn spinal cord where the pain signaling is integrated [21]. Previous report demonstrated that α3GlyRs underlies inflammatory central pain sensitization [5]. On the other hand, our results showed that both males and females had increased auxiliary βGlyR expression. This subunit is unable to form a functional channel, however, it plays an important role in the modulation of the GlyRs currents [22]. In this respect, Bormann et al., described that heteropentameric GlyR composed by α and β subunits, expressed in spinal neurons, requires a higher glycine concentration to reach about 50% of maximal effect on receptor activation in comparison to homopentameric receptors, (EC_50_ for αGlyR and αβGlyR is 30 and 54 μM, respectively) [23]. Consistent with this effect, our results showed increased α and βGlyR subunit levels after the inflammatory condition. We propose that in inflammatory pain conditions, α1βGlyRs, α2βGlyRs and α3βGlyRs could be increased to compensate for the disinhibition produced by the central sensitization process. However, in female rats the inhibitory compensation is only mediated by α3 and β subunits, suggesting that female response is insufficient to counterbalance inflammatory hypersensitivity. In agreement with these results, we showed that female rats reduced GLP-1 hormone expression after the Zymosan injection. It was reported that GLP-1 hormone had an antinociceptive effect in formalin injected male rats [11,24]. This effect was related with the modulation of opioidergic system [25]. Hence, it is possible to propose that increased female nociceptive response compared to males could be related in part with reduced spinal inhibitory compensatory mechanisms.

## Perspectives and significance

The current work contributes to the understanding of central inflammatory pain sensitization mechanisms in males and females. We have demonstrated that female rats have a lower nociceptive withdrawal threshold after inflammatory injury induced by a Zymosan A injection. We have also shown that peripheral inflammatory nociceptive sex differences is associated with differential expression levels of GlyR subunits as well as the GLP-1 hormone, suggesting that these changes could act as a compensatory mechanism to overcome central inflammatory pain sensitization.

## ACKNOWLEDGEMENTS

We thank Nicolas Sepulveda for collecting part of Western blot data and Juan Ferrada for collecting part of behavioral data.

## Contributions

GA collected and analyzed and created the figures and discussed the results. JP collected analyzed data and reviewed the manuscript. RB designed the study, collected data and discussed the results. JS designed the study, discussed the results and reviewed the manuscript. WG discussed the results and reviewed the manuscript. KC discussed the results and reviewed the manuscript. Finally, TM designed the study, collected data, discussed the results and wrote the manuscript.

